# Expanded Catalog of Gold Standard Ciliary Gene List: Integration of Novel Ciliary Genes into the Ciliome

**DOI:** 10.1101/2025.08.22.671678

**Authors:** Ferhan Yenisert, Oktay I. Kaplan

**Affiliations:** Rare Disease Laboratory, School of Life and Natural Sciences, Abdullah Gül University, Kayseri, Türkiye

## Abstract

Ciliopathies are rare genetic disorders caused by mutations in genes that encode proteins critical for cilium assembly and function, affecting various subcompartments, including the axoneme, ciliary membrane, transition zone, and basal body. These mutations lead to diverse developmental and degenerative conditions across multiple organ systems. The 2021 ciliary Gold Standard gene list (688 genes) provides a foundational resource, but many ciliary genes remain unidentified or unclassified. We developed CiliaHub (https://theciliahub.github.io/), a scalable and user-friendly platform that integrates automated literature mining with expert manual curation to identify human ciliary genes. The approach combines gene names, synonyms, and targeted keywords (e.g., cilia, flagella, basal body) to extract gene-specific mentions from published studies, followed by rigorous validation through independent expert review.

CiliaHub identified 2011 experimentally supported ciliary genes, including 1323 not previously listed in curated databases. Notably, some of these genes are co-found in both cilia and other organelles, such as mitochondria and lysosomes. Cross-referencing with mouse knockout model data revealed strong associations with ciliopathy-relevant phenotypes, such as infertility in *Cep70* and renal defects or hearing abnormalities in genes like *TMEM145*. This work also provides novel ciliopathy candidate genes for further investigation. CiliaHub provides a broad platform for fundamental and translational research, enabling the effective prioritization of candidate disease genes, including recently identified ciliopathy candidates.

## Introduction

The functional integrity of eukaryotic cells depends on the precise spatial organization and stoichiometry of proteins within subcellular organelles. Among these, cilia stand out as highly compartmentalized structures essential for sensing and transducing extracellular signals. Tissue homeostasis, embryonic development, and cellular signaling pathways such as PDGF, Wnt, and Hedgehog all depend on it^1^. The first link between ciliary dysfunction and human disease was established in 1976 when electron microscope studies revealed that ultrastructural defects in motile cilia were the cause of Primary Ciliary Dyskinesia (PCD)^2^.

The known importance of this organelle greatly increased in 2000, when abnormal ciliary signaling in renal cells was identified as the cause of cyst formation in Polycystic Kidney Disease (PKD)^3^. This was soon followed by the characterization of Bardet-Biedl Syndrome (BBS)^4^, Alström Syndrome^5^, Meckel-Gruber Syndrome^6^, Senior-Løken Syndrome^7^, Joubert Syndrome^8,9^, and Nephronophthisis (NPHP)^10^ as disorders of ciliary dysfunction, cementing the concept of a broad class of diseases known as ciliopathies comprising more than 40 different diseases^11^.

The established link between cilia and human disease has fueled a growing interest in identifying the complete proteociliome since the early 2000s. Researchers have employed a wide array of methodologies, including proteomics, bioinformatics, phylogenetic profiling, transcriptomics, high-throughput genetic screens, and patient genomics to identify ciliary components across species. These efforts have collectively yielded numerous datasets and databases (e.g., Ciliome, CiliaCarta, SysCilia Gold Standard, CentrosomeDB, CiliaMiner, and CilioGenics) with contributions from model organisms such as *Chlamydomonas, Tetrahymena, Strongylocentrotus purpuratus* (sea urchin), *Nematostella vectensis* (sea anemone), *C. elegans, Drosophila*, mouse, and human cell lines^6,11–60, 60–66^.

Despite these advances, the full extent of the human ciliary gene set remains unresolved. Many genes with clear experimental or phenotypic links to ciliary biology are still absent from curated databases. There is a growing need for a scalable and continuously updated resource that bridges automated discovery with expert validation.

To meet this challenge, we present an updated and expanded catalog of human ciliary genes through CiliaHub (https://theciliahub.github.io/), an openly accessible, literature-driven platform designed to dynamically capture and curate the growing body of knowledge in ciliary biology. CiliaHub integrates a multi-tiered approach that begins with automated text mining of the biomedical literature, utilizing gene symbols and aliases in combination with cilium-related keywords (e.g., *cilia, cilium, flagella, basal body, axoneme*, and *transition zone*). This high-throughput extraction pipeline identifies putative gene-mention relationships across thousands of PubMed-indexed publications.

CiliaHub increases the known human ciliome from 688 genes (2021 Gold Standard list) to over 2,000 curated human genes. Notably, more than 1300 of these genes are novel additions, not present in prior compilations or widely used ciliary gene databases. Many of these genes display ciliopathy-like phenotypes in animal models (e.g., hydrocephalus, situs inversus, male infertility, retinal degeneration, and renal cystic disease), which reflects their potential biological and clinical importance. Our approach contributes to the prioritizing of novel candidate genes for functional studies and clinical analysis by broadening the known ciliome.

## Material and Methods

### Data Retrieval and Literature Mining

We developed an algorithm to perform a systematic search of PubMed for the discovery of all genes related to cilia and flagella. First, we download all human gene names. Simply, a complete list of 41,078 unique human gene symbols (HGNC) (**Supplementary Table 1**) was obtained from the Ensembl database (version 110 or latest available) using the biomaRt package in R. Duplicate entries and missing values were removed to ensure data quality. To associate each gene with relevant ciliary research, a custom R function was developed to automate literature searches on PubMed. For each gene, a query combined the gene symbol with ciliary- and flagellar-related keywords: “cilia”, “cilium”, “transition zone”, “basal body”, “ciliogenesis”, “flagella”, and “flagellum”. Up to 200 of the most recent publications matching each query were retrieved.

The rvest package was employed to scrape PubMed search results programmatically. From each result, the PubMed ID (PMID) and article title were extracted. The function incorporated error handling using tryCatch to manage missing data or retrieval issues. All data were aggregated into a single data frame for downstream analysis. Software and R Environment All analyses were performed in R (version 4.4.1) using the following packages and their versions: rvest (version 1.0.4), dplyr (version 1.1.4), writexl (version 1.4.5), readxl (version 1.4.5), and rentrez (version 1.2.3).

### Protein–Protein Interaction (PPI) Network Construction

Protein–protein interaction data were retrieved using the STRING database (STRINGdb R package), which provides experimentally validated and computationally predicted interactions. Known ciliary proteins were defined based on curated databases (e.g., SYSCILIA gold standard) and literature annotations. Protein identifiers were mapped to STRING database IDs using STRINGdb functions. The get_interactions() function was used to retrieve high-confidence interactions (combined score ≥ 0.7) between known and candidate proteins. Interaction data were processed with dplyr and data.table to remove duplicates and self-interactions. An igraph object was generated for network representation, which was then converted to a tidygraph object for plotting. Known ciliary proteins were labeled red, and new ciliary proteins were labeled blue. Visualization was performed using ggraph and ggplot2, applying a force-directed layout to position nodes based on interaction strength. The node size was proportional to the degree (number of connections) within the network.

### Data Acquisition and Processing for Protein Domains

A dataset of 2,004 unique ciliary genes was curated (compiled as of August 1, 2025) using the readxl package in R (version 4.3.1). The complete set of human genes was included.

InterPro domains for both ciliary and all human genes were obtained using getBM queries to Ensembl, filtering on HGNC symbols and retrieving Ensembl gene IDs, HGNC symbols, InterPro IDs, and domain descriptions. Domains with missing or empty descriptions were excluded to ensure data quality.

### Domain Comparison and Statistical Analysis

Domain frequencies were calculated separately for ciliary genes (CiliaGeneCount) and all human genes (TotalGeneCount). A full join combined these counts, with NA values replaced by zeros. For each domain, the following metrics were computed: Non-ciliary gene count: TotalGeneCount - CiliaGeneCount. Fold enrichment: (CiliaGeneCount /2,004) /(TotalGeneCount /41,078). Fisher’s exact test for enrichment (alternative = “greater”) and depletion (alternative = “less”) using a 2×2 contingency matrix: [Cilia with domain, Non-cilia with domain; Cilia without domain, Non-cilia without domain]. P-values were adjusted using the Benjamini-Hochberg method to account for multiple testing. Significance classification: Enriched if adjusted p-value (enrichment) < 0.05 and fold enrichment > 1; Depleted if adjusted p-value (depletion) < 0.05 and fold enrichment < 1; otherwise, not significant. Log-transformed metrics: log_2_(fold enrichment) and -log_10_(adjusted p-value) for visualization purposes. Analyses were performed in R using packages tidyverse, biomaRt, ggrepel, forcats, purrr, and patchwork. Visualizations were created with ggplot2 and exported as PNG files at 300 DPI.

### Reactome pathways, KEGG biological pathways, Gene Ontology (GO)

Functional enrichment analyses were performed for human genes. First, all known and candidate genes were loaded from an Excel file, and unique and non-missing gene symbols were extracted from the “Gene” column. These gene symbols were converted to Entrez IDs using the org.Hs.eg.db database.

The enrichment analyses were performed using the ClusterProfiler and ReactomePA containers. Analyses included: (i) Reactome pathways (enrichPathway), (ii) KEGG biological pathways (enrichKEGG), (iii) Gene Ontology (GO) biological information (BP), (iv) containing (CC), and (v) functions (MF). For each analysis, p-value improvements were observed, and highly significant pathways were trimmed (**Supplementary Figure**).

The analysis results were tabulated, and dotplot visualizations were generated using ggplot2 using fold enrichment and -log10 (adj p-value) values for the top 20 most significant pathways. In addition, Reactome pathway-gene mappings were generated, and the number of different pathways involved for each gene was calculated. All results were compiled into an Excel file, providing detailed reports of both pathway information and gene-pathway relationships. Analyses were conducted using the R programming language and relevant bioinformatics packages.

### Computational Human Ciliopathy Phenotype Analysis

The Open Targets platform was used for the analysis of human genes and associated ciliopathy phenotypes. Only the first Ensembl ID of each gene was selected from the gene list prepared for analysis^67^. The Open Targets GraphQL API was used to query associated diseases for each gene, and associations with a score ≥ 0.5 were considered.

Phenotypes associated with ciliopathy were identified using literature and clinical data sources. The associations between these phenotypes and genes were compared to create a binary matrix (presence/absence). The phenotype matrix was converted to long-form, and dot plots were created to visualize the gene-phenotype associations.

Furthermore, to systematically examine gene-phenotype associations, bipartite networks were created. Phenotypes and genes were represented as nodes, and colors were used to distinguish phenotype types. All analyses were performed using R software and the ggplot2 and ggraph packages.

### Computational Mouse Ciliopathy Phenotype Analysis

Phenotypic effects of mouse genes were analyzed using R (≥ 4. x). Analysis was performed using the packages readxl, dplyr, tidyr, stringr, httr, jsonlite, writexl, and ggplot2. The input gene list was read from an Excel or CSV file, and the names were normalized (first letter uppercase, all others lowercase). Phenotype data for each gene were queried via International Mouse Phenotyping Consortium (IMPC) REST API^68^. Results from all pages were combined, and duplicate phenotypes were removed to create a unique list. Ciliopathy-associated phenotypes were mapped to predefined keywords, and if a match occurred, the gene was marked with “Ciliopathy_Flag = YES.” Phenotypes were mapped into a matrix with TRUE/FALSE values, with phenotype terms as rows and genes as columns. Outputs were recorded as a per-gene phenotype list and a phenotype × gene matrix, with the genes with the most phenotypes visualized using a dot plot. Analyses were designed to be reproducible with modular R scripts.

### Comparison of MitoCarta, CiliaHub, and Lyosome

For our comparative analysis, we utilized three primary gene sets: MitoCarta, CiliaHub, and a predictive lysosome gene list. The MitoCarta gene list (https://www.broadinstitute.org/mitocarta/mitocarta30-inventory-mammalian-mitochondrial-proteins-and-pathways) and the predictive lysosome gene list were downloaded on August 13, 2025. The lysosome gene set, specifically the “LOCATE Predicted Protein Localization Annotations,” was obtained from the Ma’ayan Lab’s Harmonizome database (https://maayanlab.cloud/Harmonizome/gene_set/lysosome/LOCATE+Predicted+Protein+Localization+Annotations).

### Data Analysis and Figures

All algorithms, data processing, network analyses, and figure generation codes for this study were managed through our CiliaHub platform. The complete source code and associated data are publicly available on GitHub, ensuring full reproducibility of our results. CiliaHub on GitHub: https://github.com/thekaplanlab/CiliaHub

## Results

### Development and Curation of the CiliaHub Gene Resource

We developed an algorithm called CiliaHub that applies cilia-specific keyword filtering (cilia, cilium, flagella, basal body, axoneme, transition zone) to PubMed data, systematically screening 24,482 protein-coding genes to identify ciliary and cilia-associated genes. Gene inclusion was determined by two criteria: (i) experimental localization evidence (genes with confirmed localization to cilia, designated as cilia-localizing), and (ii) functional evidence (genes lacking direct localization data but linked to ciliary function, designated as cilia-associated).

The initial search yielded 5917 likely ciliary and cilia-associated genes. From this pool, generated primarily through automated filtering, we confirmed 1382 human genes as newly recognized ciliary genes, encompassing both cilia-localizing and cilia-associated categories (**Figure 1**). For instance, although *SMC1A* and *SMC3* were reported as cilia-localizing as early as 2005, they had not been formally included in prior ciliary gene lists^69^. Likewise, *ITGB1* (integrin β1) and *CD44* were shown to localize to cilia in 2020 but were also absent from prior compilations^70^.

**Figure 1:**
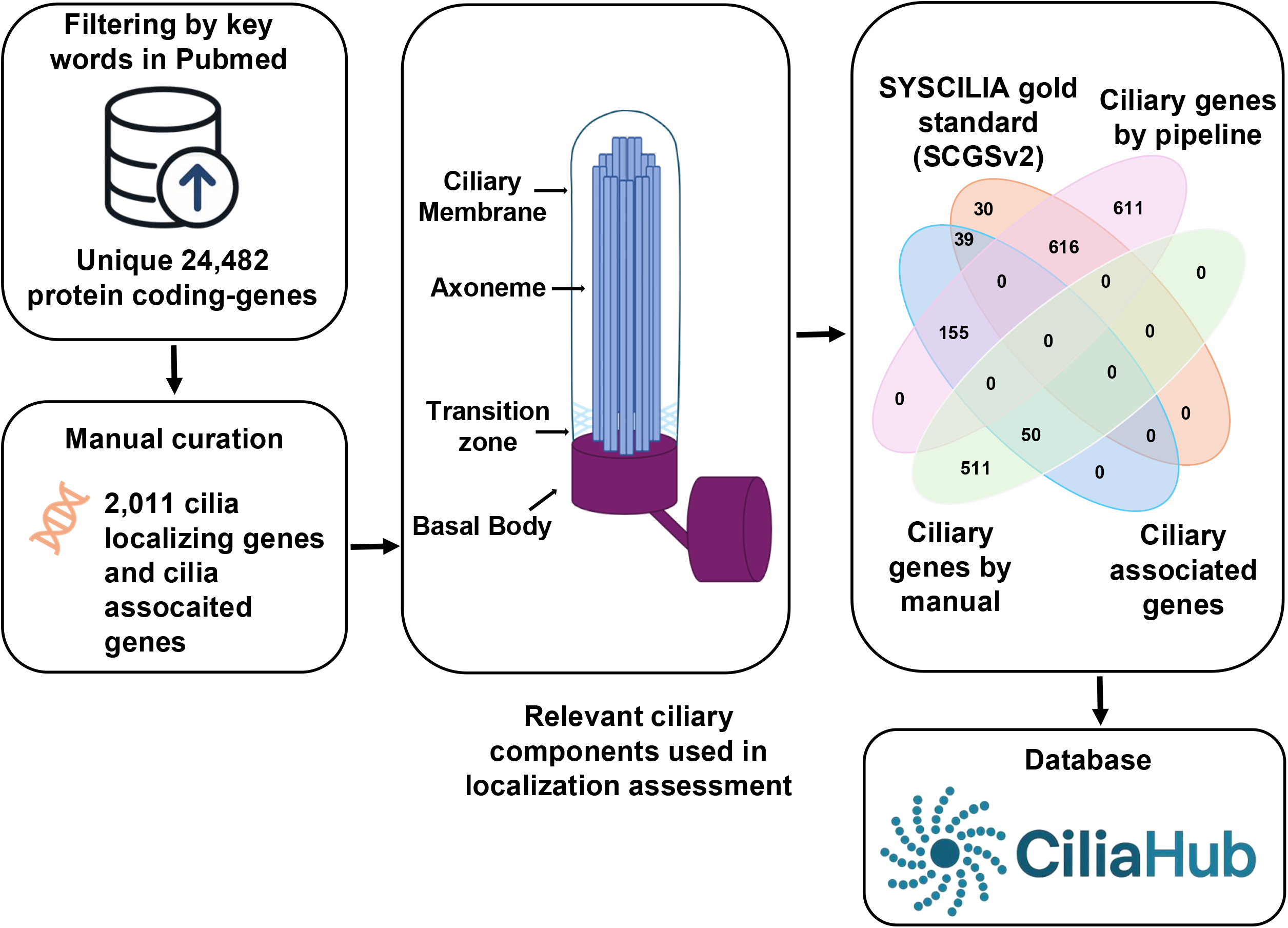
CiliaHub Curation Pipeline and Gene Discovery. This figure illustrates the multi-step process used to develop the CiliaHub gene resource. The pipeline begins with an extensive literature search of the PubMed database using a combination of gene names, synonyms, and specific ciliary keywords (e.g., cilia, cilium, flagella, basal body, axoneme, and transition zone). This automated filtering step generated an initial pool of 5900 candidate genes. This was followed by a rigorous manual curation process, where an expert reviewed the literature for each candidate to confirm its association with ciliary structures or function. This manual review resulted in a refined list of 2011 ciliary and cilia-associated genes. The figure shows the relationship between the CiliaHub list and the previous SYSCILIA Gold Standard (SCGSv2) list, highlighting the overlap and the significant number of novel genes (1323) identified by our platform. These novel genes are a key output of CiliaHub, providing important opportunities for the ciliopathy field.

We compared our dataset against the SYSCILIA Gold Standard (SCGSv2), which contains 688 curated ciliary genes^40,54^. With our analysis, the number of known ciliary genes nearly doubled. Subsequent manual curation expanded the dataset further, encompassing both cilia-localizing and cilia-associated genes, and increasing the total to 2,011 genes (**Supplementary Table 2A, B, C and, D**). Ultimately, CiliaHub provides a unified and expanded resource that significantly improves the coverage of ciliary genes beyond existing databases. To facilitate access, we developed a publicly available CiliaHub website (https://theciliahub.github.io/), where users can explore and download the curated gene lists.

### Domain Enrichment in ciliary genes

The analysis included 2,004 ciliary and cilia-associated genes, representing approximately 4.9% of the 41,078 unique HGNC symbols retrieved from the Ensembl database (**Figure 2A and Supplementary Tables 1 and 2A**). A total of 3945 unique InterPro domains were identified in ciliary genes, comprising 21 % of the 18,663 unique domains found across all human genes (**Figure 2A**). Statistical analysis identified 392 domains that are considerably enriched in ciliary genes (adjusted p < 0.05, fold enrichment > 1), with the top 20 showing fold enrichments ranging from approximately 3.3 to 19.6 (**Figure 2B**). The volcano plot illustrated domain significance, with enriched domains in the positive log_2_(fold enrichment) quadrant above the - log_10_(p) = 1.3 threshold (p = 0.05) and depleted domains in the negative quadrant. Dotted lines at log_2_(fold enrichment) = ±1 marked moderate effect sizes, while non-significant domains clustered near zero enrichment (**Figure 2C**). Notable enriched domains included “Coiled-coil domain,” “Armadillo-type fold,” “Protein kinase-like domain superfamily,” “Kinesin motor domain,” and “Tubulin/FtsZ families,” which are critical for ciliary assembly, intracellular transport, and microtubule organization (**Supplementary Table 3**). In contrast, 5 domains were substantially depleted in ciliary genes (adjusted p < 0.05, fold enrichment < 1). The top depleted domains, absent from all ciliary genes (CiliaGeneCount = 0), were prevalent in non-ciliary genes, such as “Immunoglobulin-like receptors” in immune regulation and “Krueppel C2H2-type zinc-finger,” appearing in hundreds of non-ciliary genes (Supplementary Table 3). These results highlight the distinct domain composition of ciliary genes, enriched in structural and motility-related domains while depleted in regulatory and immune-related domains, reflecting their specialized biological roles.

**Figure 2:**
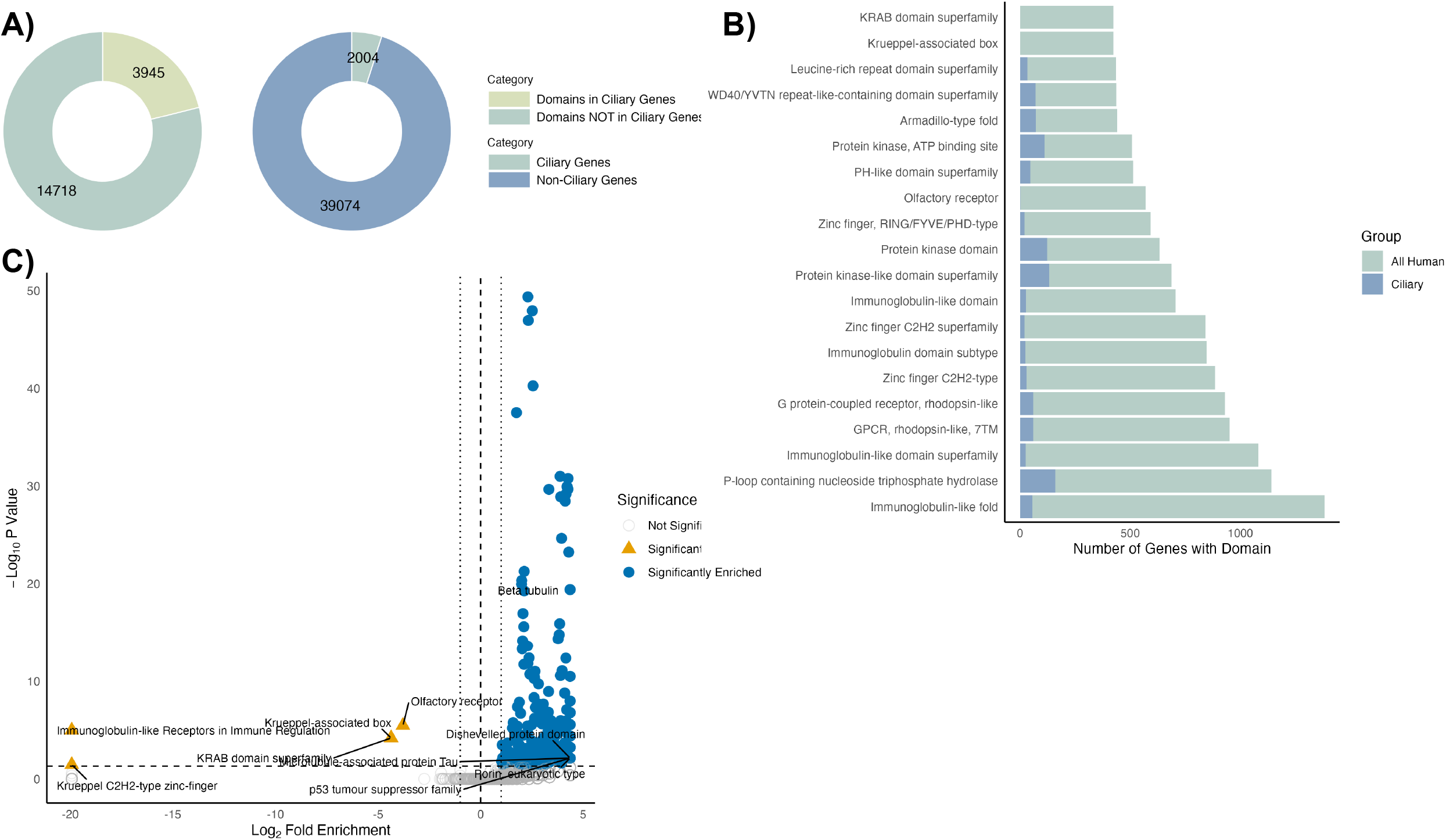
Analysis of Ciliary and Non-Ciliary Gene Domain Enrichment. **(A)** The pie charts illustrate the distribution of genes based on their association with cilia. The chart on the left shows the total number of genes in the CiliaHub dataset, categorized into those with domains found in ciliary genes (light green) and those with domains not found in ciliary genes (dark green). The chart on the right shows the same data for the entire human genome, highlighting the relatively small but highly significant fraction of genes with ciliary-related functions. **(B)** This bar chart shows the protein domains (Top 20 Most Enriched Protein Domains in Ciliary Genes) that are most frequently found within the CiliaHub ciliary gene list compared to the overall human genome. Each bar represents a specific protein domain, with its length corresponding to the number of genes containing that domain. The two colors distinguish between the domain counts in all human genes (light blue) and specifically in ciliary genes (dark blue), providing a visual representation of how enriched these domains are in ciliary proteins. **(C)** The volcano plot visualizes the statistical significance and fold-enrichment of protein domains in ciliary genes. The x-axis represents the Log2 Fold Enrichment, indicating how much more frequently a domain appears in ciliary genes compared to non-ciliary genes. The y-axis represents the -Log10 P-Value, with higher values indicating a more statistically significant enrichment. Each dot represents a protein domain. Domains that are enriched in ciliary genes are shown in blue, while domains with no enrichment are in yellow or orange, highlighting a small number of domains that are underrepresented.

### Analysis of Specific Gene Interactions in Ciliary PPI Network

We next examined the protein–protein interaction (PPI) network between known ciliary genes and newly identified candidates. The network forms a highly interconnected structure. In the primary network (**Figure 3A and Supplementary Table 4**), blue nodes represent known ciliary genes, and red nodes indicate newly identified ones. The central cluster contains numerous nodes and edges, suggesting that some new genes participate in established ciliary processes by interacting with well-studied proteins.

**Figure 3:**
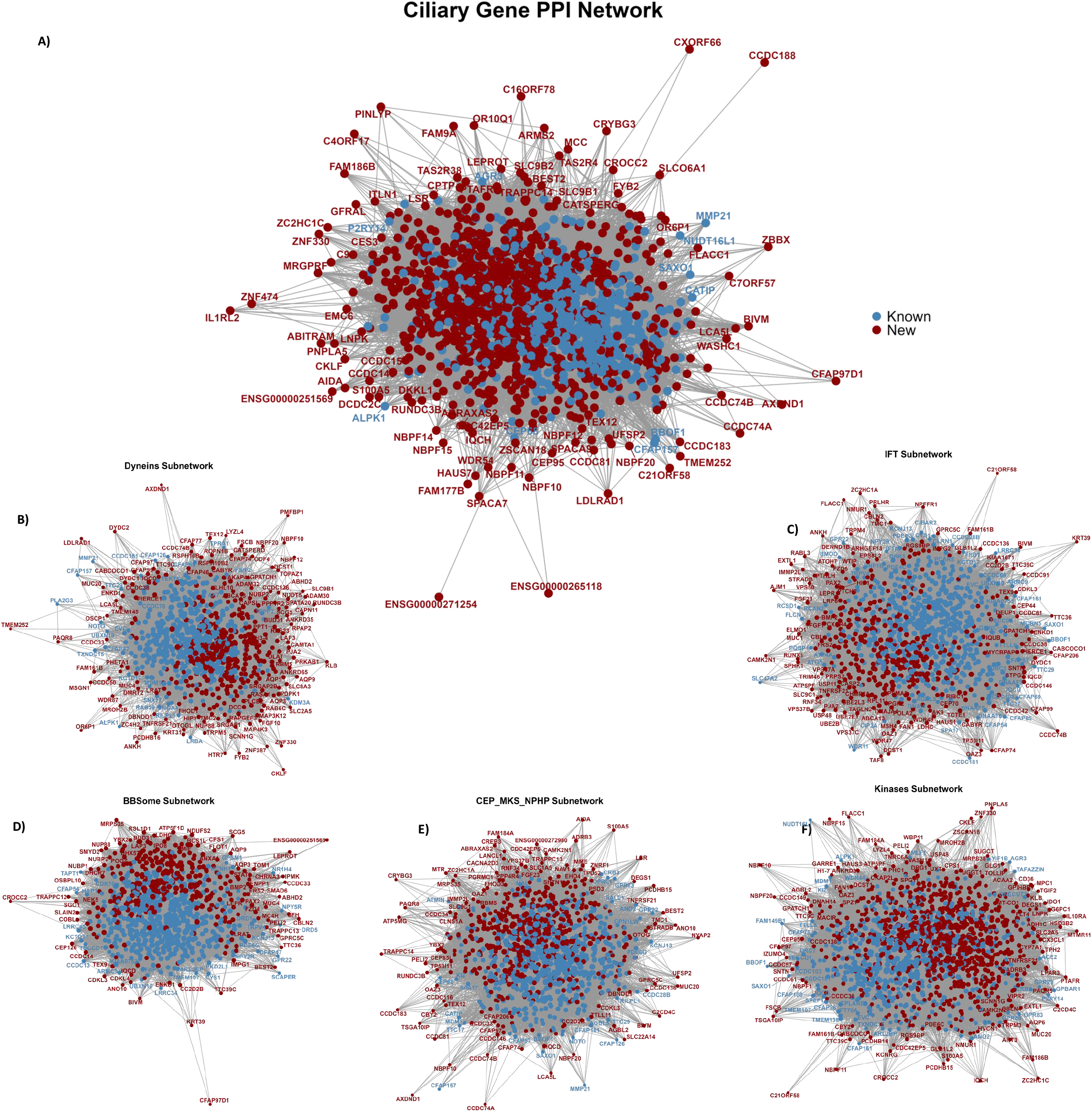
Protein–protein interaction (PPI) network of known and novel ciliary proteins. The network displays experimentally supported and predicted interactions between ciliary proteins (blue nodes) and newly identified candidate ciliary proteins (orange nodes). Six major subnetworks are highlighted: **(A)** Ciliary gene core PPI network, **(B)** Dynein-associated subnetwork, **(C)** BBSome complex subnetwork, **(D)** CEP-MKS-NPHP transition zone subnetwork, **(E)** Kinase-enriched subnetwork, and **(F)** other associated signaling proteins. Node labels represent gene symbols; bold text indicates proteins validated in this study, and asterisks denote proteins with previously uncharacterized ciliary roles. Edges represent protein–protein associations from curated databases, experimental data, and computational predictions. Known ciliary proteins are derived from curated ciliary gene datasets, while novel candidates were identified from screening and functional assays.

Singly isolated red nodes, often located at the network periphery, likely represent underexplored ciliary components. Their limited connectivity may reflect either a lack of experimental interaction data or genuinely novel roles within the ciliary system. For example, CCDC188, shown as a peripheral red node, has minimal interactions, reinforcing its status as an undercharacterized gene. In contrast, CEP164 is a known ciliary gene that shows extensive connections with other established ciliary genes such as MKS1 and RPGRIP1L, consistent with its well-defined role at the transition zone (**Figure 3B**).

Subnetwork analysis reveals distinct functional modules. The dynein subnetwork, representing the retrograde IFT motor, interacts greatly with known ciliary proteins, reflecting its critical role in regulating ciliary structure and motility. Similarly, the IFT, BBSome, and MKS–NPHP subnetworks display dense connectivity, reflecting their importance in ciliary trafficking and structural organization (**Figure 3C, D, and E**).

The Kinome subnetwork (**Figure 3F**) illustrates the regulatory role of kinases within ciliary biology. Phosphorylation functions as a molecular switch, modulating protein activity, stability, and interactions. Kinase–ciliary protein interactions suggest diverse regulatory mechanisms: for example, phosphorylation of IFT proteins could alter transport speed or cargo binding, while phosphorylation of structural components may trigger ciliary disassembly. Mapping these kinase-centered PPIs may provide critical insight into the signaling pathways controlling ciliary assembly, disassembly, and function. Dysregulation of the kinome–cilia network may contribute to ciliopathies such as Bardet–Biedl syndrome, polycystic kidney disease, and retinal degeneration.

### Prioritization of Candidate Genes for Ciliopathy

Ciliopathies encompass a diverse group of rare diseases arising from defects in ciliary structure or function, affecting single or multiple organ systems. To prioritize candidate genes implicated in cilia-associated symptoms, we exploited a curated ciliary gene list and integrated data from OpenTarget and the Mouse Phenotype Database to map gene-phenotype associations, with a focus on ciliary symptoms/phenotypes, including sinusitis, hydrocephalus, coloboma, neurological, renal, skeletal, auditory, and metabolic abnormalities.

Our analysis revealed that 77 genes were associated with ciliopathy-related phenotypes in humans (**Figure 4A and Supplementary Table S5**). Notably, genes such as *AKT3, PTEN*, and *MPDZ* were linked to neurological defects like hydrocephalus, seizures, and ataxia, while *BICC1, PKHD1L1*, and *WBP11* were associated with renal/cystic diseases, showing overlap with mouse models of kidney dysplasia (**Figure 4B and C**). Abnormal heart morphology was observed in the following genes *Ankmy2, Anxa1, Arl16, Atg7, Bicc1, Cacna2d2, Canx, Casp8, Cd36, Cobl, Crocc2, Crx, Dbn1, Dync2i1, Enpp1, Epas1, Fam161b, Fhod3, Krt34, Pebp1, Ppp1r35, Psen2, Ptprm, Rnf34, Slc9b1, Spz1, Tdo2*. Auditory dysfunction was tied to genes like *OTOG, TMC1*, and *CCDC50*, with mouse orthologs confirming hearing loss, and skeletal ciliopathies, including *BMP2* and *PIK3CD*, correlated with polydactyly and craniofacial defects.

**Figure 4.**
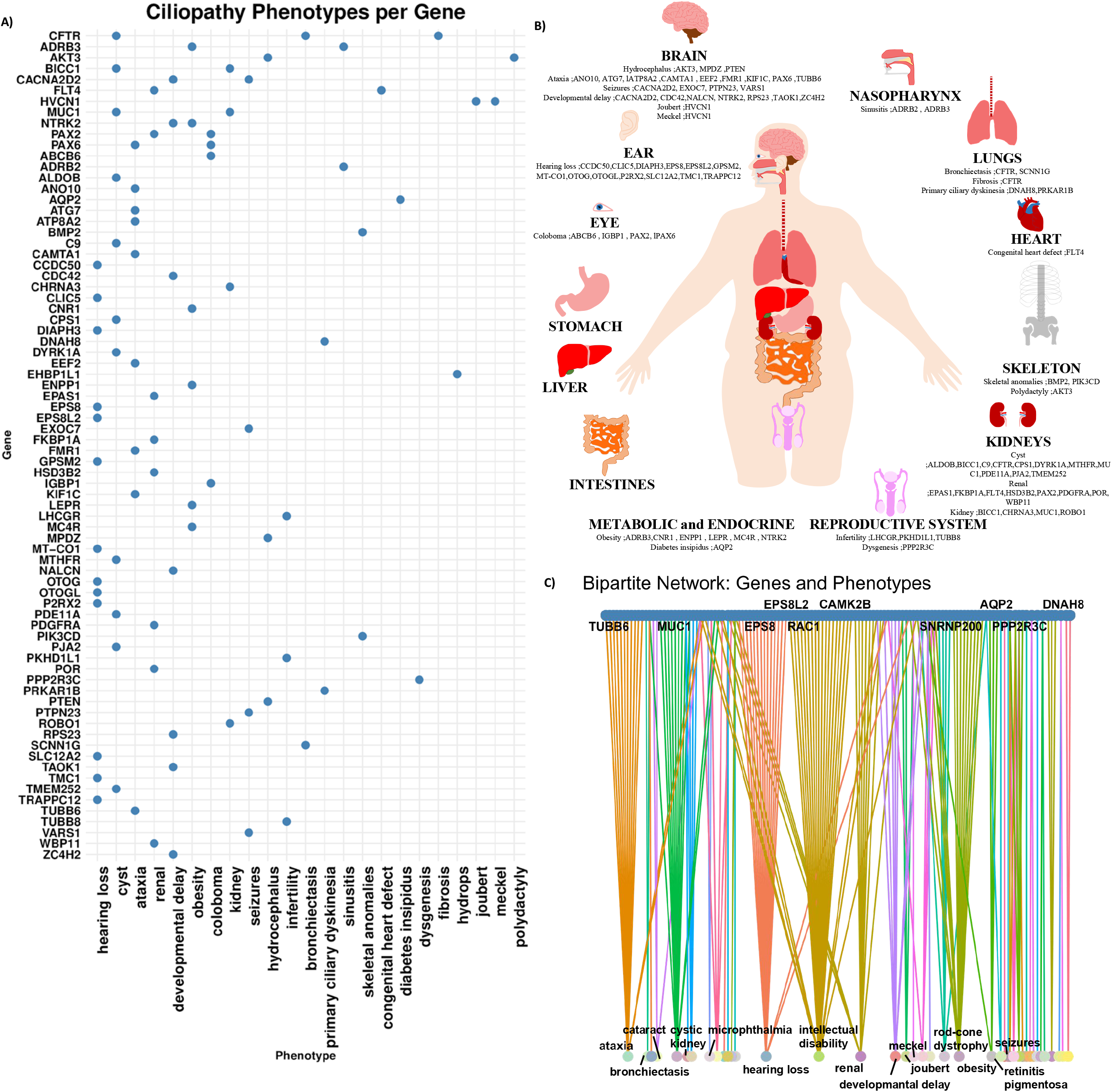
Ciliopathy phenotypes associated with individual genes in humans. **(A)** Dot plot showing the distribution of ciliopathy-related phenotypes across individual genes retrieved from OpenTarget (a score ≥ 0.5). Each dot represents an observed association between a specific gene (y-axis) and a phenotype (x-axis), highlighting the pleiotropic effects of ciliary genes. **(B)** Schematic representation of human organ systems affected by ciliopathy genes. Selected genes associated with phenotypes in each organ are indicated. This visualization demonstrates the multisystem impact of ciliary dysfunction, including the brain, ear, eye, nasopharynx, lungs, heart, skeleton, liver, kidneys, intestines, reproductive system, and metabolic/endocrine organs. **(C)** Bipartite network connecting genes (top nodes) to their associated phenotypes (bottom nodes). Edges indicate documented associations between individual genes and specific ciliopathy phenotypes. The network shows both highly pleiotropic genes, which connect to multiple phenotypes, and more specialized genes linked to specific phenotypes.

In mouse models, 113 genes (new ciliary genes) were linked to cilia-associated phenotypes, revealing additional insights not fully captured in human data, such as embryonic lethality in *Cdc42* and *Uggt1* knockouts, implicating their role in early development, infertility in *Catsper2* and *Lepr* mutants, suggesting ciliary roles in reproduction, and metabolic dysregulation in *Lepr*-associated obesity, highlighting systemic ciliary functions. Comparative analysis showed concordant phenotypes for genes like *PTEN* and *OTOG* across species (**Figure 5A, B, and Supplementary Table S5**).

**Figure 5:**
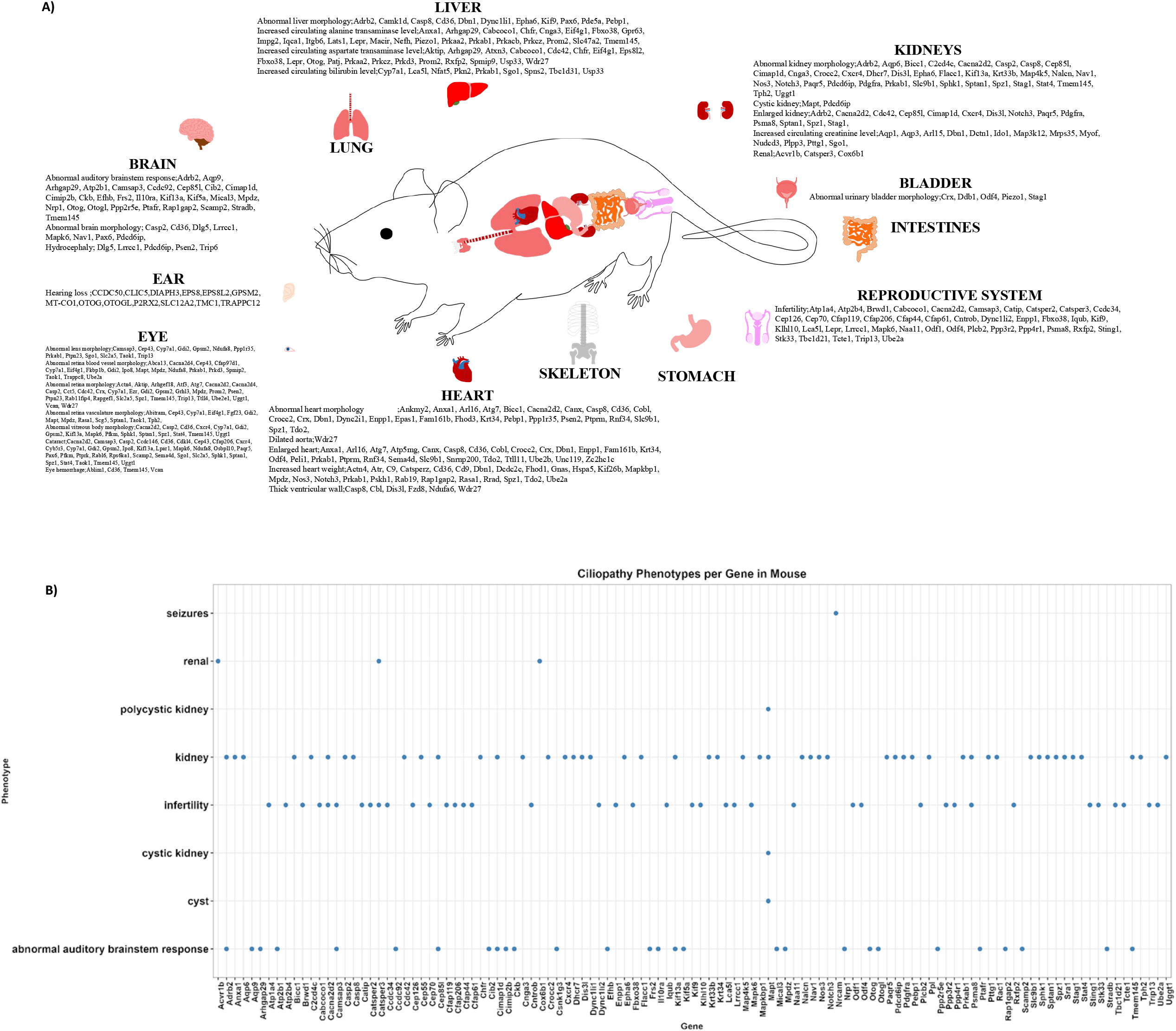
Pleiotropic Effects of Ciliopathy-Associated Genes in Mouse Models. **(A) Organ-Specific Phenotypes of Ciliopathy Genes** A schematic diagram of a mouse depicts the various organ systems affected by ciliopathy-related gene mutations. For each organ system, including the liver, brain, lung, ear, eye, skeleton, heart, stomach, intestines, reproductive system, bladder, and kidneys, a list of associated genes and the resulting abnormal phenotypes is provided. This panel visually represents the widespread impact of these genes across the body. **(B) Ciliopathy Phenotype and Gene Correlation Matrix** A dot plot mapping specific ciliopathy-related phenotypes (y-axis) to a list of known ciliopathy genes (x-axis). Each blue dot at the intersection of a gene and a phenotype indicates that a mutation in that particular gene has been experimentally shown to cause the corresponding phenotype in mouse models. This plot provides a clear overview of the phenotypic spectrum for each gene and demonstrates the significant overlap in phenotypes caused by different genes, a common feature of ciliopathies.

Overall, our prioritization strategy, combining a curated ciliary gene set with human and mouse phenotypic data, enables the identification of high-confidence candidate genes for further functional studies in ciliopathy.

## Discussion

The development of CiliaHub marks an important step in ciliary biology, increasing the catalog of known ciliary genes. Through its innovative combination of automated literature mining and expert manual curation, the platform has identified 1,314 novel ciliary genes, surpassing the 688 genes in the previous SYSCILIA Gold Standard, and bringing the total to over 2,000 genes (*Supplementary Table S2*). This dramatic expansion not only demonstrates the power of systematic curation approaches but also reveals how many ciliary-associated genes were previously overlooked in conventional databases.

The role of cilia in metabolism has been recognized. For example, *ASNS* (Asparagine synthetase) is a ciliary protein that has been identified as a key enzyme that allows primary cilia to sense the availability of the amino acid glutamine^71^. Additionally, *MTHFR* (Methylenetetrahydrofolate Reductase) is a basal body protein involved in folate and homocysteine metabolism. Maybe, disruptions in *MTHFR* function, as seen in folate metabolism disorders, may impair ciliary signaling, potentially contributing to ciliopathies like neural tube defects or polycystic kidney disease^72–74^. Furthermore, *ACLY* (ATP citrate lyase) is a ciliary-localizing metabolic enzyme that converts citrate to acetyl-CoA, a key precursor for lipid biosynthesis and protein acetylation. Importantly, loss of *ACLY* resulted in a reduction in ciliated cell differentiation^75,76^.

Analysis of the protein-protein interaction network reveals intriguing patterns in ciliary gene organization. While well-established hub genes like CEP164 show extensive interactions with known ciliary proteins, validating the reliability of the curation pipeline, the identification of peripheral nodes such as *CCDC188* and *ZNF474* is particularly exciting. These minimally connected genes likely represent specialized components with context-dependent functions that may have been missed in previous studies. The network analysis also successfully reconstructs known functional modules, including the dynein complexes, BBSome, and MKS-NPHP networks, demonstrating how novel genes integrate into established ciliary pathways. This systems-level view provides critical insights into how newly identified genes may contribute to ciliary structure and function.

Cilia do not function in isolation; their activity depends on dynamic interactions with other organelles. Each organelle performs distinct functions, but they must communicate and exchange metabolites, signals, or stress responses to stay alive and adapt. Membrane contact sites between organelles allow for direct exchange of lipids, ions, and reactive species, often bypassing vesicular trafficking pathways and enabling rapid functional coordination. For example, reactive oxygen species (ROS) transfer at peroxisome–mitochondria contact sites regulates mitochondrial redox homeostasis^77^. The peroxisomal protein ACBD5 and the mitochondrial protein PTPIP51 were shown to form physical contacts between the two organelles.

Similarly, cilia require a continuous and substantial supply of ATP to support intraflagellar transport (IFT) and other energy-demanding processes. However, cilia themselves do not contain mitochondria, the primary producers of cellular ATP. Instead, mitochondria are frequently positioned in close proximity to the ciliary base, enabling rapid delivery of ATP and potentially other metabolites to sustain ciliary activity. This spatial organization underscores the importance of physical and functional connections between cilia and mitochondria. The physiological relevance of this connection is highlighted by patient data^78^. Individuals with mitochondrial disease displayed ciliary defects that disrupted left–right patterning, leading to heterotaxy, a condition characterized by abnormal organ positioning. Notably, mitochondrial disorders and ciliopathies share overlapping clinical manifestations, including retinal dystrophy, hearing loss, diabetes, and cardiomyopathy, underscoring their potential mechanistic connections. In these patients, mitochondrial DNA content was reduced, and experimental manipulation of mitochondrial function in zebrafish, human fibroblasts, and Tetrahymena revealed direct effects on ciliogenesis. These findings establish mitochondrial abundance and function as key determinants of ciliary structure and activity.

An important question is how these cilia–mitochondria interactions are achieved at the molecular level. One possibility is that specific proteins function in both organelles, serving as molecular bridges or regulators of shared pathways such as energy metabolism, redox balance, and signaling. For example, NEK10 localizes to cilia and mitochondria and may facilitate interactions between these organelles^79,80^. Furthermore, there are many genes encoding such dual-localized or dual-function proteins, including *ARL2, CEP89, NME3, PRKACA, VDAC3*, and *ABCB6* (**Supplementary Table S6**). Further studies on these proteins could clarify the mechanisms of cilia–mitochondria coordination.

Similar functional relationships are likely to exist between cilia and lysosomes. Lysosomes serve as central hubs for cellular waste clearance, autophagy, and nutrient signaling (e.g., mTOR), all of which are processes known to regulate ciliogenesis and ciliary signaling^81,82^. Consistently, a number of genes appear to localize and/or function in both organelles (**Supplementary Table S7**). Clinical observations also point to overlap between these systems: Fabry disease, a lysosomal storage disorder, can present with kidney cysts, while polydactyly has been reported in other lysosomal diseases such as mucopolysaccharidosis, phenotypes traditionally associated with ciliopathies^11,83,84^. This link is reinforced by direct experimental evidence from mouse models of Niemann-Pick C1 disease, which demonstrate that this lysosomal disorder causes distinct alterations in the number, length, and morphology of primary cilia^85^. Taken together, these examples show that cilia do not operate in isolation but depend on dynamic interactions with neighboring organelles.

Domain enrichment patterns further support the biological relevance of the expanded ciliome. The strong representation of coiled-coil, kinesin motor, and tubulin domains aligns perfectly with the structural and mechanical requirements of cilia, while the notable absence of domains like immunoglobulin-like folds emphasizes the specialized nature of ciliary proteins. Particularly interesting is the identification of kinase-enriched subnetworks, which suggests underappreciated roles for phosphorylation-based regulation in ciliary biology. DYF-5/MAK/CILK1 phosphorylates the tubulin-binding module of IFT74 to promote tubulin unloading at the ciliary tip, thereby regulating ciliary length and morphology in *C. elegans*^86,87^. The conservation of these regulatory mechanisms, as seen in the DYF-5/MAK-IFT74 pathway in *C. elegans*, points to fundamental principles of ciliary control that may be disrupted in various ciliopathies.

The translational potential of CiliaHub becomes evident when examining the phenotypic associations of both established and novel ciliary genes. High-confidence candidates like *AKT3* and *PTEN* show remarkable conservation of neurological and renal phenotypes between human and mouse models, while the identification of 113 novel genes with cilia-associated mouse phenotypes provides opportunities for further investigation. Genes like *Catsper2* and *Lepr*, associated with infertility and metabolic dysregulation, respectively, exemplify how this resource can uncover unexpected connections between ciliary function and systemic physiology. For complex multisystem disorders like Bardet-Biedl syndrome, this expanded gene set provides new candidates for functional validation and potential therapeutic targeting.

While CiliaHub represents a major advance, certain limitations must be acknowledged. The reliance on published literature means that genes with subtle or context-specific roles may still be missing, and the manual curation process, while rigorous, may introduce some subjectivity. Future developments could incorporate machine learning approaches to improve efficiency and consistency, along with the integration of multi-omics data to capture genes with indirect roles in ciliary biology. Experimental validation will be particularly important for assessing predicted protein interactions and the functional significance of novel candidates.

CiliaHub provides an unprecedented resource that expands the known ciliome and supports scientists in accelerating discoveries into the mechanisms of human disease. The platform is publicly accessible at https://rarediseaselab.github.io/home/#ciliahub. By bridging high-throughput data mining with expert curation, the platform offers both breadth and reliability, serving as a foundation for future discoveries in ciliary biology. The publicly accessible interface ensures that this resource will be valuable to both basic researchers studying ciliary mechanisms and clinicians investigating ciliopathies. As the field progresses, CiliaHub’s dynamic curation approach positions it to incorporate new findings and continue expanding our knowledge of this crucial cellular organelle and its associated disorders.

## Acknowledgment

The authors gratefully acknowledge financial support from the Health Institutes of Türkiye (TÜSEB) under grant number 44276 to O.I.K. We also thank DoÜancan Özbek for assistance with interactive visualization of the expression analysis on the website.

## Data Sharing

All data can be downloaded from the https://theciliahub.github.io/.

## Notes

https://theciliahub.github.io/

### Competing Interest Statement

The authors have declared no competing interest.

## References

1. Satir, P. & Christensen, S. T. Overview of structure and function of mammalian cilia. Annu. Rev. Physiol. 69, 377–400 (2007).

2. Afzelius, B. A. A human syndrome caused by immotile cilia. Science 193, 317–319 (1976).

3. Pazour, G. J. et al. Chlamydomonas IFT88 and its mouse homologue, polycystic kidney disease gene tg737, are required for assembly of cilia and flagella. J. Cell Biol. 151, 709–718 (2000).

4. Ansley, S. J. et al. Basal body dysfunction is a likely cause of pleiotropic Bardet-Biedl syndrome. Nature 425, 628–633 (2003).

5. Bahl, S. et al. Evidence of a common founder for SCA12 in the Indian population. Ann. Hum. Genet. 69, 528–534 (2005).

6. Smith, J. C., Northey, J. G. B., Garg, J., Pearlman, R. E. & Siu, K. W. M. Robust method for proteome analysis by MS/MS using an entire translated genome: demonstration on the ciliome of Tetrahymena thermophila. J. Proteome Res. 4, 909–919 (2005).

7. Otto, E. A. et al. Nephrocystin-5, a ciliary IQ domain protein, is mutated in Senior-Loken syndrome and interacts with RPGR and calmodulin. Nat. Genet. 37, 282–288 (2005).

8. Valente, E. M. et al. Mutations in CEP290, which encodes a centrosomal protein, cause pleiotropic forms of Joubert syndrome. Nat. Genet. 38, 623–625 (2006).

9. Bielas, S. L. et al. Mutations in INPP5E, encoding inositol polyphosphate-5-phosphatase E, link phosphatidyl inositol signaling to the ciliopathies. Nat. Genet. 41, 1032–1036 (2009).

10. Otto, E. A. et al. Mutations in INVS encoding inversin cause nephronophthisis type 2, linking renal cystic disease to the function of primary cilia and left-right axis determination. Nat. Genet. 34, 413–420 (2003).

11. Turan, M. G., Orhan, M. E., Cevik, S. & Kaplan, O. I. CiliaMiner: an integrated database for ciliopathy genes and ciliopathies. Database J. Biol. Databases Curation 2023, baad047 (2023).

12. Ostrowski, L. E. et al. A proteomic analysis of human cilia: identification of novel components. Mol. Cell. Proteomics MCP 1, 451–465 (2002).

13. Avidor-Reiss, T. et al. Decoding cilia function: defining specialized genes required for compartmentalized cilia biogenesis. Cell 117, 527–539 (2004).

14. Blacque, O. E. et al. Functional Genomics of the Cilium, a Sensory Organelle. Curr. Biol. 15, 935–941 (2005).

15. Pazour, G. J., Agrin, N., Leszyk, J. & Witman, G. B. Proteomic analysis of a eukaryotic cilium. J. Cell Biol. 170, 103–113 (2005).

16. Keller, L. C., Romijn, E. P., Zamora, I., Yates, J. R. & Marshall, W. F. Proteomic analysis of isolated chlamydomonas centrioles reveals orthologs of ciliary-disease genes. Curr. Biol. CB 15, 1090–1098 (2005).

17. Inglis, P. N., Boroevich, K. A. & Leroux, M. R. Piecing together a ciliome. Trends Genet. TIG 22, 491–500 (2006).

18. Chen, N. et al. Identification of ciliary and ciliopathy genes in Caenorhabditis elegans through comparative genomics. Genome Biol. 7, R126 (2006).

19. Kilburn, C. L. et al. New Tetrahymena basal body protein components identify basal body domain structure. J. Cell Biol. 178, 905–912 (2007).

20. Liu, Q. et al. The proteome of the mouse photoreceptor sensory cilium complex. Mol. Cell. Proteomics MCP 6, 1299–1317 (2007).

21. Li, J. B. et al. Comparative genomics identifies a flagellar and basal body proteome that includes the BBS5 human disease gene. Cell 117, 541–552 (2004).

22. Efimenko, E. et al. Analysis of xbx genes in C. elegans. Dev. Camb. Engl. 132, 1923–1934 (2005).

23. Laurençon, A. et al. Identification of novel regulatory factor X (RFX) target genes by comparative genomics in Drosophila species. Genome Biol. 8, R195 (2007).

24. Mayer, U. et al. The proteome of rat olfactory sensory cilia. Proteomics 9, 322–334 (2009).

25. Mayer, U. et al. Proteomic analysis of a membrane preparation from rat olfactory sensory cilia. Chem. Senses 33, 145–162 (2008).

26. Ross, A. J., Dailey, L. A., Brighton, L. E. & Devlin, R. B. Transcriptional profiling of mucociliary differentiation in human airway epithelial cells. Am. J. Respir. Cell Mol. Biol. 37, 169–185 (2007).

27. McClintock, T. S., Glasser, C. E., Bose, S. C. & Bergman, D. A. Tissue expression patterns identify mouse cilia genes. Physiol. Genomics 32, 198–206 (2008).

28. Stubbs, J. L., Oishi, I., Izpisúa Belmonte, J. C. & Kintner, C. The forkhead protein Foxj1 specifies node-like cilia in Xenopus and zebrafish embryos. Nat. Genet. 40, 1454–1460 (2008).

29. Nogales-Cadenas, R., Abascal, F., Díez-Pérez, J., Carazo, J. M. & Pascual-Montano, A. CentrosomeDB: a human centrosomal proteins database. Nucleic Acids Res. 37, D175–180 (2009).

30. Fritz-Laylin, L. K. & Cande, W. Z. Ancestral centriole and flagella proteins identified by analysis of Naegleria differentiation. J. Cell Sci. 123, 4024–4031 (2010).

31. Müller, H. et al. Proteomic and functional analysis of the mitotic Drosophila centrosome. EMBO J. 29, 3344–3357 (2010).

32. Kim, J. et al. Functional genomic screen for modulators of ciliogenesis and cilium length. Nature 464, 1048–1051 (2010).

33. Jakobsen, L. et al. Novel asymmetrically localizing components of human centrosomes identified by complementary proteomics methods. EMBO J. 30, 1520–1535 (2011).

34. Geremek, M. et al. Gene expression studies in cells from primary ciliary dyskinesia patients identify 208 potential ciliary genes. Hum. Genet. 129, 283–293 (2011).

35. Phirke, P. et al. Transcriptional profiling of C. elegans DAF-19 uncovers a ciliary base-associated protein and a CDK/CCRK/LF2p-related kinase required for intraflagellar transport. Dev. Biol. 357, 235–247 (2011).

36. Lauwaet, T. et al. Mining the Giardia genome and proteome for conserved and unique basal body proteins. Int. J. Parasitol. 41, 1079–1092 (2011).

37. Ivliev, A. E., ‘t Hoen, P. A. C., van Roon-Mom, W. M. C., Peters, D. J. M. & Sergeeva, M. G. Exploring the transcriptome of ciliated cells using in silico dissection of human tissues. PloS One 7, e35618 (2012).

38. Hoh, R. A., Stowe, T. R., Turk, E. & Stearns, T. Transcriptional program of ciliated epithelial cells reveals new cilium and centrosome components and links to human disease. PloS One 7, e52166 (2012).

39. Ishikawa, H., Thompson, J., Yates, J. R. & Marshall, W. F. Proteomic analysis of mammalian primary cilia. Curr. Biol. CB 22, 414–419 (2012).

40. van Dam, T. J. et al. The SYSCILIA gold standard (SCGSv1) of known ciliary components and its applications within a systems biology consortium. Cilia 2, 7 (2013).

41. van Dam, T. J. P. et al. CiliaCarta: An integrated and validated compendium of ciliary genes. PloS One 14, e0216705 (2019).

42. Choksi, S. P., Babu, D., Lau, D., Yu, X. & Roy, S. Systematic discovery of novel ciliary genes through functional genomics in the zebrafish. Dev. Camb. Engl. 141, 3410–3419 (2014).

43. Chung, M.-I. et al. Coordinated genomic control of ciliogenesis and cell movement by RFX2. eLife 3, e01439 (2014).

44. Roosing, S. et al. Mutations in CEP120 cause Joubert syndrome as well as complex ciliopathy phenotypes. J. Med. Genet. 53, 608–615 (2016).

45. Wheway, G. et al. An siRNA-based functional genomics screen for the identification of regulators of ciliogenesis and ciliopathy genes. Nat. Cell Biol. 17, 1074–1087 (2015).

46. Mick, D. U. et al. Proteomics of Primary Cilia by Proximity Labeling. Dev. Cell 35, 497–512 (2015).

47. Lambacher, N. J. et al. TMEM107 recruits ciliopathy proteins to subdomains of the ciliary transition zone and causes Joubert syndrome. Nat. Cell Biol. 18, 122–131 (2016).

48. Boldt, K. et al. An organelle-specific protein landscape identifies novel diseases and molecular mechanisms. Nat. Commun. 7, 11491 (2016).

49. Shaheen, R. et al. Characterizing the morbid genome of ciliopathies. Genome Biol. 17, 242 (2016).

50. Shim, H. et al. Function-driven discovery of disease genes in zebrafish using an integrated genomics big data resource. Nucleic Acids Res. 44, 9611–9623 (2016).

51. Sigg, M. A. et al. Evolutionary Proteomics Uncovers Ancient Associations of Cilia with Signaling Pathways. Dev. Cell 43, 744–762.e11 (2017).

52. Nevers, Y. et al. Insights into Ciliary Genes and Evolution from Multi-Level Phylogenetic Profiling. Mol. Biol. Evol. 34, 2016–2034 (2017).

53. Breslow, D. K. et al. A CRISPR-based screen for Hedgehog signaling provides insights into ciliary function and ciliopathies. Nat. Genet. 50, 460–471 (2018).

54. Vasquez, S. S. V., van Dam, J. & Wheway, G. An updated SYSCILIA gold standard (SCGSv2) of known ciliary genes, revealing the vast progress that has been made in the cilia research field. Mol. Biol. Cell 32, br13 (2021).

55. May, E. A. et al. Time-resolved proteomics profiling of the ciliary Hedgehog response. J. Cell Biol. 220, e202007207 (2021).

56. Rao, V. G. & Kulkarni, S. S. Xenopus to the rescue: A model to validate and characterize candidate ciliopathy genes. Genes. N. Y. N 2000 59, e23414 (2021).

57. Pir, M. S. et al. CilioGenics: an integrated method and database for predicting novel ciliary genes. Nucleic Acids Res. 52, 8127–8145 (2024).

58. Dobbelaere, J., Su, T. Y., Erdi, B., Schleiffer, A. & Dammermann, A. A phylogenetic profiling approach identifies novel ciliogenesis genes in Drosophila and C. elegans. EMBO J. 42, e113616 (2023).

59. Van Sciver, R. E. & Caspary, T. A prioritization tool for cilia-associated genes and their in vivo resources unveils new avenues for ciliopathy research. Dis. Model. Mech. 17, dmm052000 (2024).

60. Lindskog, C. et al. A high-resolution spatial map of cilia-associated proteins based on characterization of the human fallopian tube-specific proteome. Preprint at 10.21203/rs.3.rs-3914234/v1 (2024).

61. Fredrick, A. S. et al. Identifying Ciliary Proteins in Mammalian Retinas using a Gentle Extraction Method. MicroPublication Biol. 2024, (2024).

62. Lange, S. M., Eisert, R. J. & Brown, A. A conserved mechanism for the retrieval of polyubiquitinated proteins from cilia. 2025.04.24.650332 Preprint at 10.1101/2025.04.24.650332 (2025).

63. Aarts, E. M. et al. Network-based framework for studying etiology and phenotype diversity in primary ciliopathies. 2025.01.08.631887 Preprint at 10.1101/2025.01.08.631887 (2025).

64. Liang, C., Wang, W. & Chen, J. First transcriptome assembly of a new ciliate species (Protocruzia marianaensis) isolated from the Mariana Trench area. Mar. Genomics 79, 101164 (2025).

65. Hansen, J. N. et al. Intrinsic Heterogeneity of Primary Cilia Revealed Through Spatial Proteomics. 2024.10.20.619273 Preprint at 10.1101/2024.10.20.619273 (2025).

66. Macarelli, V. et al. Proximity proteomics of primary cilia in human hypothalamic neurons. 2025.05.11.653368 Preprint at 10.1101/2025.05.11.653368 (2025).

67. Carvalho-Silva, D. et al. Open Targets Platform: new developments and updates two years on. Nucleic Acids Res. 47, D1056–D1065 (2019).

68. Muñoz-Fuentes, V. et al. The International Mouse Phenotyping Consortium (IMPC): a functional catalogue of the mammalian genome that informs conservation. Conserv. Genet. Print 19, 995–1005 (2018).

69. Khanna, H. et al. RPGR-ORF15, which is mutated in retinitis pigmentosa, associates with SMC1, SMC3, and microtubule transport proteins. J. Biol. Chem. 280, 33580–33587 (2005).

70. Lee, M. N. et al. The primary cilium directs osteopontin-induced migration of mesenchymal stem cells by regulating CD44 signaling and Cdc42 activation. Stem Cell Res. 45, 101799 (2020).

71. Steidl, M. E. et al. Primary cilia sense glutamine availability and respond via asparagine synthetase. Nat. Metab. 5, 385–397 (2023).

72. Toriyama, M., Toriyama, M., Wallingford, J. B. & Finnell, R. H. Folate-dependent methylation of septins governs ciliogenesis during neural tube closure. FASEB J. Off. Publ. Fed. Am. Soc. Exp. Biol. 31, 3622–3635 (2017).

73. van der Put, N. M. et al. Mutated methylenetetrahydrofolate reductase as a risk factor for spina bifida. Lancet Lond. Engl. 346, 1070–1071 (1995).

74. Shillingford, J. M., Leamon, C. P., Vlahov, I. R. & Weimbs, T. Folate-conjugated rapamycin slows progression of polycystic kidney disease. J. Am. Soc. Nephrol. JASN 23, 1674–1681 (2012).

75. Kim, B. R. et al. The oxygen level in air directs airway epithelial cell differentiation by controlling mitochondrial citrate export. Sci. Adv. 11, eadr2282 (2025).

76. Hatzivassiliou, G. et al. ATP citrate lyase inhibition can suppress tumor cell growth. Cancer Cell 8, 311–321 (2005).

77. DiGiovanni, L. F. et al. ROS transfer at peroxisome-mitochondria contact regulates mitochondrial redox. Science 389, 157–162 (2025).

78. Burkhalter, M. D. et al. Imbalanced mitochondrial function provokes heterotaxy via aberrant ciliogenesis. J. Clin. Invest. 129, 2841–2855 (2019).

79. Porpora, M. et al. Counterregulation of cAMP-directed kinase activities controls ciliogenesis. Nat. Commun. 9, 1224 (2018).

80. Peres de Oliveira, A. et al. NEK10 interactome and depletion reveal new roles in mitochondria. Proteome Sci. 18, 4 (2020).

81. Morleo, M. et al. Crosstalk between cilia and autophagy: implication for human diseases. Autophagy 19, 24–43 (2023).

82. Prosseda, P. P., Dannewitz Prosseda, S., Tran, M., Liton, P. B. & Sun, Y. Crosstalk between the mTOR pathway and primary cilia in human diseases. Curr. Top. Dev. Biol. 155, 1–37 (2023).

83. Gupta, S. et al. Temporal expression profiling identifies pathways mediating effect of causal variant on phenotype. PLoS Genet. 11, e1005195 (2015).

84. Mehta, A., Beck, M., Linhart, A., Sunder-Plassmann, G. & Widmer, U. History of lysosomal storage diseases: an overview. in Fabry Disease: Perspectives from 5 Years of FOS (eds Mehta, A., Beck, M. & Sunder-Plassmann, G.) (Oxford PharmaGenesis, Oxford, 2006).

85. Lucarelli, M. et al. Anomalies in Dopamine Transporter Expression and Primary Cilium Distribution in the Dorsal Striatum of a Mouse Model of Niemann-Pick C1 Disease. Front. Cell. Neurosci. 13, 226 (2019).

86. Jiang, X. et al. DYF-5/MAK-dependent phosphorylation promotes ciliary tubulin unloading. Proc. Natl. Acad. Sci. U. S. A. 119, e2207134119 (2022).

87. Sezer, A. et al. A homozygous frameshift variant in the CILK1 gene causes cranioectodermal dysplasia. Eur. J. Hum. Genet. EJHG (2025) doi:10.1038/s41431-025-01902-0.

